# Efficient installation of heterozygous mutations in human pluripotent stem cells using prime editing

**DOI:** 10.1101/2025.05.25.656043

**Authors:** Annabelle Suter, Alison Graham, Jia Yi Kuah, Jason Crisologo, Chathuni Gunatilake, Koula Sourris, Michael See, Fernando J Rossello, Mirana Ramialison, Katerina Vlahos, Sara E Howden

**Affiliations:** Murdoch Children’s Research Institute, The Royal Children’s Hospital, Melbourne, VIC, Australia; Novo Nordisk Foundation Center for Stem Cell Medicine, Murdoch Children’s Research Institute, Melbourne, Victoria, Australia; Department of Paediatrics, University of Melbourne, Melbourne, Victoria, Australia; Department of Clinical Pathology, University of Melbourne, Melbourne, VIC, Australia; Australian Regenerative Medicine Institute, Monash University, Victoria, Australia

## Abstract

The utility of human pluripotent stem cells (hPSCs) is greatly enhanced by the ability to introduce precise, site-specific genetic modifications with minimal off-target effects. Although Cas9 endonuclease is an exceptionally efficient gene-editing tool, its propensity for generating biallelic modifications often limits its capacity for introducing heterozygous variants. Here, we use prime editing (PE) to install heterozygous edits in over ten distinct genetic loci, achieving knock-in efficiencies of up to 40% without the need for subsequent purification or drug selection steps. Moreover, PE enables the precise introduction of heterozygous edits in paralogous genes that are otherwise extremely challenging to achieve using endonuclease-based editing approaches. We also show that PE can be successfully combined with reprogramming to derive heterozygous iPSC clones directly from human fibroblasts and peripheral blood mononuclear cells. Our findings highlight the utility of PE for generating hPSCs with complex edits and represents a powerful platform for modelling disease-associated dominant mutations and gene-dosage effects in an entirely isogenic context.

## INTRODUCTION

Human induced pluripotent stem cells (iPSCs) hold incredible promise for many exciting applications, including autologous cell-based therapies, disease modelling, and drug discovery. However, harnessing the full potential of iPSCs depends largely on an ability to genetically modify cells in a manner that is both highly precise and efficient. In the context of disease modelling, this may entail correcting mutations in patient-derived iPSCs or introducing specific genetic variants into an otherwise healthy iPSC line. Indeed, the introduction of specific genetic variants into a well-characterised hPSC clone enables its functional consequence to be studied in a completely isogenic context, thereby eliminating the substantial variability that can arise from genetic background differences. As an example, we recently described the introduction a series of homozygous mutations in the *NPHS2* locus in wildtype iPSCs to reveal a previously unappreciated reduction in variant protein (PODOCIN), variant-specific subcellular localisation, as well as specific effects on association with another important glomerular protein, NEPHRIN.^1^

While technologies such as CRISPR-Cas9 have revolutionized genome engineering workflows and offer the ability to edit the genome with high precision and efficacy,^2-4^ achieving heterozygous modifications - where only one allele carries the desired edit - remains challenging. This can be attributed to the remarkable efficiency of CRISPR/Cas9 in generating DNA double-stranded breaks, which typically results in cells containing the intended (knock-in) or unintended (indel) edit within both alleles of a given target locus.

Here, we explore the use of prime-editing (PE) to reliably and efficiently install heterozygous edits in iPSCs without impacting the second allele. The PE system is comprised of a Cas9-nickase fused to a reverse transcriptase (nCas9-RT) complexed to a programmable PE guide RNA (pegRNA).^5^ The pegRNA functions both to direct nCas9-RT and, once reverse transcribed, as a repair template at the target site. Compared to genome editing methods that rely on DNA double-stranded breaks, PE provides a significantly higher ratio of successful edits to unintended insertions or deletions (indels) and is also thought to be less dependent on endogenous repair mechanisms. PE has also been shown to exhibit minimal off-target effects^6^, further strengthening its appeal in genome engineering applications. While successful application of PE in hPSCs has been previously described^7-10^, we extend this work further to show that PE is an effective strategy for generating hPSCs with “clean” heterozygous edits in a broad range of genetic loci, including within genes that harbour identical sequences within closely related paralogues. We also demonstrate that PE can be combined with episomal reprogramming to derive iPSC clones with heterozygous edits directly from skin fibroblasts or patient blood samples. Collectively, these findings highlight the utility of PE as an effective strategy for generating iPSC clones with complex genetic modifications.

## RESULTS

Truncating mutations in *TTN* are one of the leading causes of dilated cardiomyopathy.^11^ We initially assessed the ability of PE to introduce a heterozygous *TTN* c.72663delA mutation in the wildtype iPSC line, PB010.5 (Figure S1A). A pegRNA comprising a *TTN*-specific spacer, 12 bp primer binding site (PBS) and 22 bp reverse transcription template (RTT) encoding the truncating *TTN* c.72663delA mutation and a nearby synonymous change (*TTN* c.72657T>A) (Figure S1B) was incorporated into a plasmid-based vector for expression from a U6 promoter. The evopreQ_1_ motif was also included, downstream of the RTT, to enhance pegRNA stability as previously described.^12^ The resulting epegRNA plasmid was introduced into iPSCs via electroporation using our previously optimised workflow^13^, along with mRNA encoding either the PEmax^14^ or more recently described PE6b^15^ prime editors. We also evaluated the effect of a secondary downstream “nicking” sgRNA specific to the complementary strand as described in the PE3 (and PE5) system^14^, and/or expression of the engineered dominant-negative MLH1dn mutant as described in the PE4 and PE5 systems.^14,16^ Gene editing outcomes were evaluated in the transfected bulk cell population 2-3 days after electroporation (Figure 1A) and in isolated colonies (Figure 1B) by amplicon next generation sequencing (NGS). To attain independent iPSC clones harbouring heterozygous knock-in of the *TTN* c.72663delA mutation, we selected colonies/wells with a *TTN* c.72663delA allele frequency >10% but ≤50% for subcloniong. At least ten subclones were then isolated and genotyped by NGS to attain purely heterozygous subclones, harbouring a 1:1 ratio of *TTN* c.72663delA to wildtype reads. An overview of the workflow used throughout this study is depicted in Figure S2.

**Figure 1.**
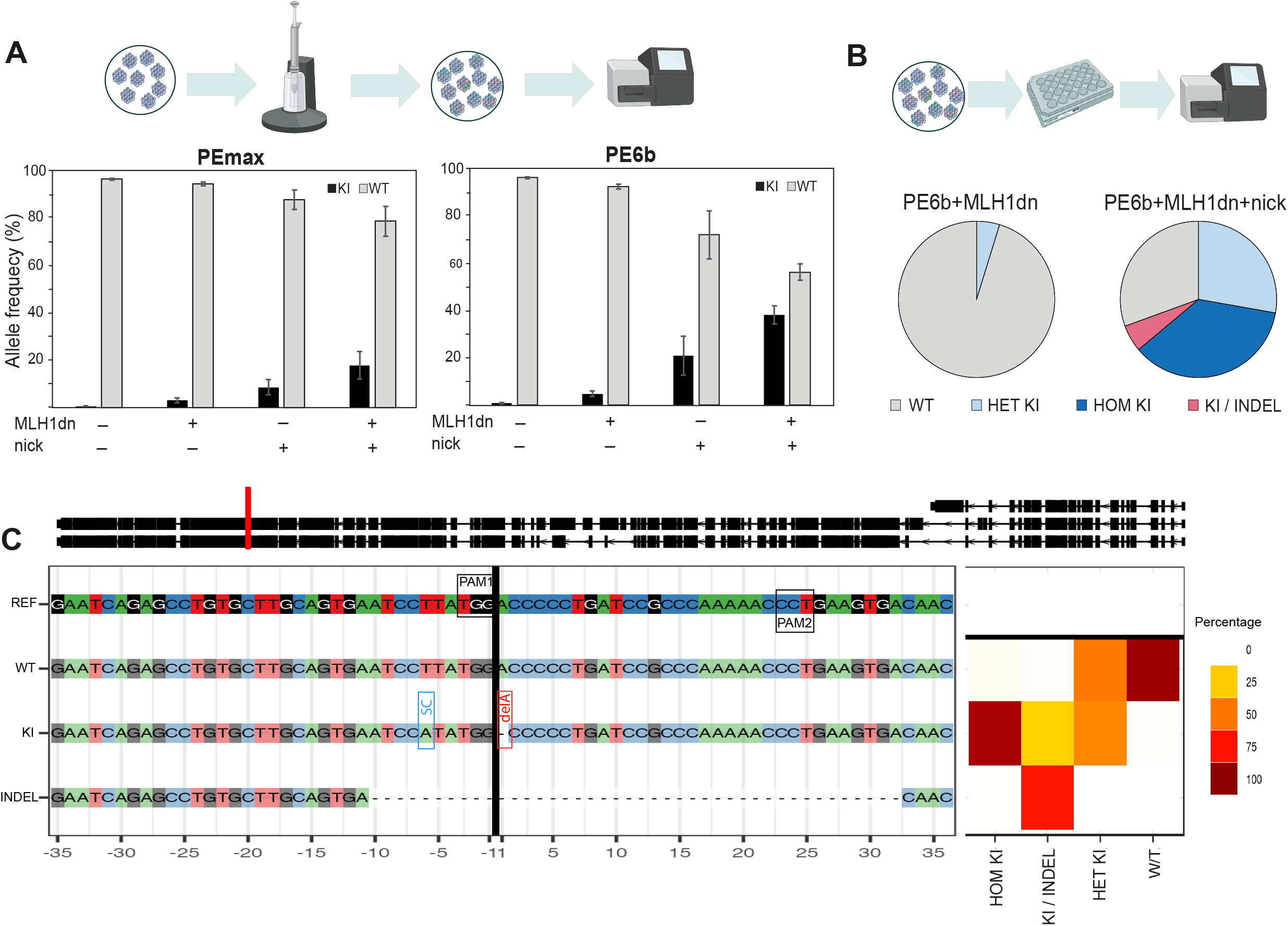
Evaluation of prime editing strategies in iPSCs. (A) Evaluation of knock-in efficiency, measured as *TTN* c.72663delA allele frequency, in bulk cultures following introduction of PE factors in healthy iPSCs. N=3. Error bars indicate standard deviation (SD), N=number of biological replicates. (B) Genotypic analysis of individual colonies isolated from PE6b + MHL1dn transfection experiments performed with and without the nicking sgRNA. (C) CrispRVariants plot of representative iPSC clones harbouring homozygous and heterozygous knock-in (with and without indel) of the *TTN* c.72663delA mutation and accompanying synonymous change (*TTN* c.72657T>A), attained from the “PE6b + MHL1dn + nick” condition.

Compared to PEmax, PEb6 resulted in a consistent albeit modest improvement in editing efficiency, as detected by NGS analysis of bulk-transfected cultures (Figure 1A). While the inclusion of either MLH1dn mRNA or the nicking sgRNA alone enhanced editing efficiency, the most dramatic increase occurred when both were used together, resulting in ∼40% of total alleles harbouring the *TTN* c.72663delA mutation (Figure 1A; PE6b + MLH1dn + nick condition). We next performed NGS analysis on isolated colonies, with a focus on the PE6b + MLH1dn ± nick conditions (Figure 1B). In the PE6b + MHLdn1 condition, of 42 colonies screened we identified two (5%) heterozygous for the *TTN* c.72663delA mutation, which was confirmed following subcloning analysis (Figure S2B). Notably, of the 36 colonies screened from the PE6b + MHLdn1 + nick condition, the *TTN* c.72663delA variant was detected in 25 (69%) colonies; of which 10 were likely heterozygous (≤50% *TTN* c.72663delA alleles), 13 homozygous (>50% *TTN* c.72663delA alleles) and two harboured heterozygous knock-in but also a deletion spanning the intervening sequence between the PE and nicking protospacers (Figure 1C).

We also conducted a direct comparison of editing outcomes using a typical homology-directed repair (HDR) strategy. An RNP complex consisting of Cas9 nuclease and the same *TTN*-specific sgRNA sequence encoded by the epegRNA, along with a single-stranded oligonucleotide (ssODN) encoding the *TTN* c.72663delA mutation and *TTN* c.72657T>A synonymous change (Figure S1C) was introduced into MCRIi010-A iPSCs, with gene editing outcomes assessed as described above. NGS analysis of the bulk transfected population revealed knock-in efficiencies comparable to that attained for the best performing PE condition, with ∼42% of total alleles carrying both the *TTN* c.72663delA mutation and synonymous change (Figure S3A). We also detected a high frequency of “partial knock-in” (∼11% of total alleles), where incorporation of the synonymous change occurred in the absence of the *TTN* c.72663delA mutation. Such partial knock-in events were far less common across all tested PE conditions, with the highest incidence (∼1% of total alleles) observed in the PE6b + MLH1dn + nick condition (Figure S3B and S3E). As expected, we also observed a much higher incidence of indel mutations (∼16% of total alleles) in the RNP condition (Figure S3D) compared to all PE conditions. Indels were barely detectable (≤0.1% of total reads) in the absence of the nicking sgRNA and remained low (∼1% of total reads) in PE conditions that included the nicking sgRNA (Figure S3E). NGS analysis of individual colonies isolated from the RNP condition revealed a broad spectrum of genotypes, with complete homozygous knock-in of the *TTN* c.72663delA mutation and accompanying synonymous change being the most prevalent outcome (∼40% of total colonies screened). Reflective of the bulk analysis, indel mutations were also common, detected in more than a third (∼36%) of iPSC colonies screened. Notably, heterozygous knock-in of the *TTN* c.72663delA mutation (in the absence of any secondary indel mutations) was only detected in two colonies, although both were homozygous for the *TTN* c.72657T>A synonymous change, indicating knock-in events on both alleles (one being partial).

We further assessed the capacity of PE to introduce heterozygous single nucleotide variants (SNV) in an additional six genetic loci (*FND3CB, SFTPA2, MYBPC3, RTEL1, INF2, RAG1*) across four additional iPSC lines (Table 1). We again performed a head-to-head comparison with PE6b and PEmax prime editors for installation of the *TTN* c.72663delA mutation but in the female IPSC line (MCRIi005-A), as well for two additional SNVs (*FNDC3B* c.2206G>A and *SFTPA2* c.160C>T) in the commercially available ChiPSC18 line (Cellartis/Takara Bio). In agreement with our previous findings targeting the *TTN* locus in MCRIi010-A iPSCs, PE6b consistently resulted in a modest increase in editing efficiency compared to PEmax. We also evaluated the effect of a secondary nick in six separate editing experiments (*MYBPC3* c.1928-2A>G, *RTEL1* c.3791G>A, *RTEL1* c.2920C>T, *RAG1* c.108C>T, *INF2 c*.599T>G and *INF2 c*.643A>C) and generally observed pronounced increases in editing efficiency in experiments that included a nicking sgRNA. Indeed, for two SNVs (*MYBPC3* c.1928-2A>G and *RTEL1* c.2920C>T), the addition of a nicking sgRNA was essential, with knock-in reads undetectable in bulk transfected cultures where the nicking sgRNA was omitted (Table 1). While we observed a broad variation in editing efficiency (ranging from 0.3–30.5% in bulk cultures) we successfully derived at least one, but typically multiple, independent iPSC clones harbouring heterozygous incorporation of each of the SNVs tested (Figure 2). Although several iPSC clones carrying indel mutations were also identified, these were predominantly observed in PE conditions that included a secondary nicking sgRNA and were typically associated with knock-in of the gene-specific SNV (Figure 2).

**Table 1.** Summary of PE experiments performed across four different healthy iPSC lines, targeting seven genetic loci. Targeting (knock-in) efficiency was measured by amplicon NGS in bulk transfected cultures.

| **SNV** | **Cell name** | **PEmax / PE6b** | **Nick** | **KI allele frequency (% total reads)** |
| --- | --- | --- | --- | --- |
| *TTN* c.72663delA | PB005 (hPSCreg:MCRIi005-A) | PEmax | No | 1.97 |
|  |  | PE6b | No | 2.13 |
| *FNDC3B* c.2206G>A | ChiPSC18 (Cellartis) | PEmax | No | 1.69 |
|  |  | PE6b | No | 2.09 + |
| *SFTPA2* c.160C>T | ChiPSC18 (Cellartis) | PEmax | No | 4.45 |
|  |  | PE6b | No | 4.84 + |
| *MYBPC3* c.1928-2A>G | PB010 (hPSCreg:MCRIi010-A) | PE6b | No | 0 |
|  |  | PE6b | Yes | 0.30 + |
| *RTEL1* c.3791G>A | ChiPSC18 (Cellartis) | PE6b | No | 3.21 + |
|  |  | PE6b | Yes | 30.50 + |
| *RTEL1* c.2920C>T | ChiPSC18 (Cellartis) | PE6b | No | 0 |
|  |  | PE6b | Yes | 8.35 + |
| *RAG1* c.108 C>T | PB001 (hPSCreg:MCRIi001-A) | PE6b | No | 1.57 |
|  |  | PE6b | Yes | 13.07 + |
| *INF2* c.599 T>G | PB001 (hPSCreg:MCRIi001-A) | PE6b | No | 0.54 |
|  |  | PE6b | Yes | 9.83 |
| *INF2* c.643 A>C | PB001 (hPSCreg:MCRIi001-A) | PE6b | No | 7.43 + |
|  |  | PE6b | Yes | 8.01 |

+ indicates experiments where individual colonies were isolated and genotyped by NGS.

**Figure 2.**
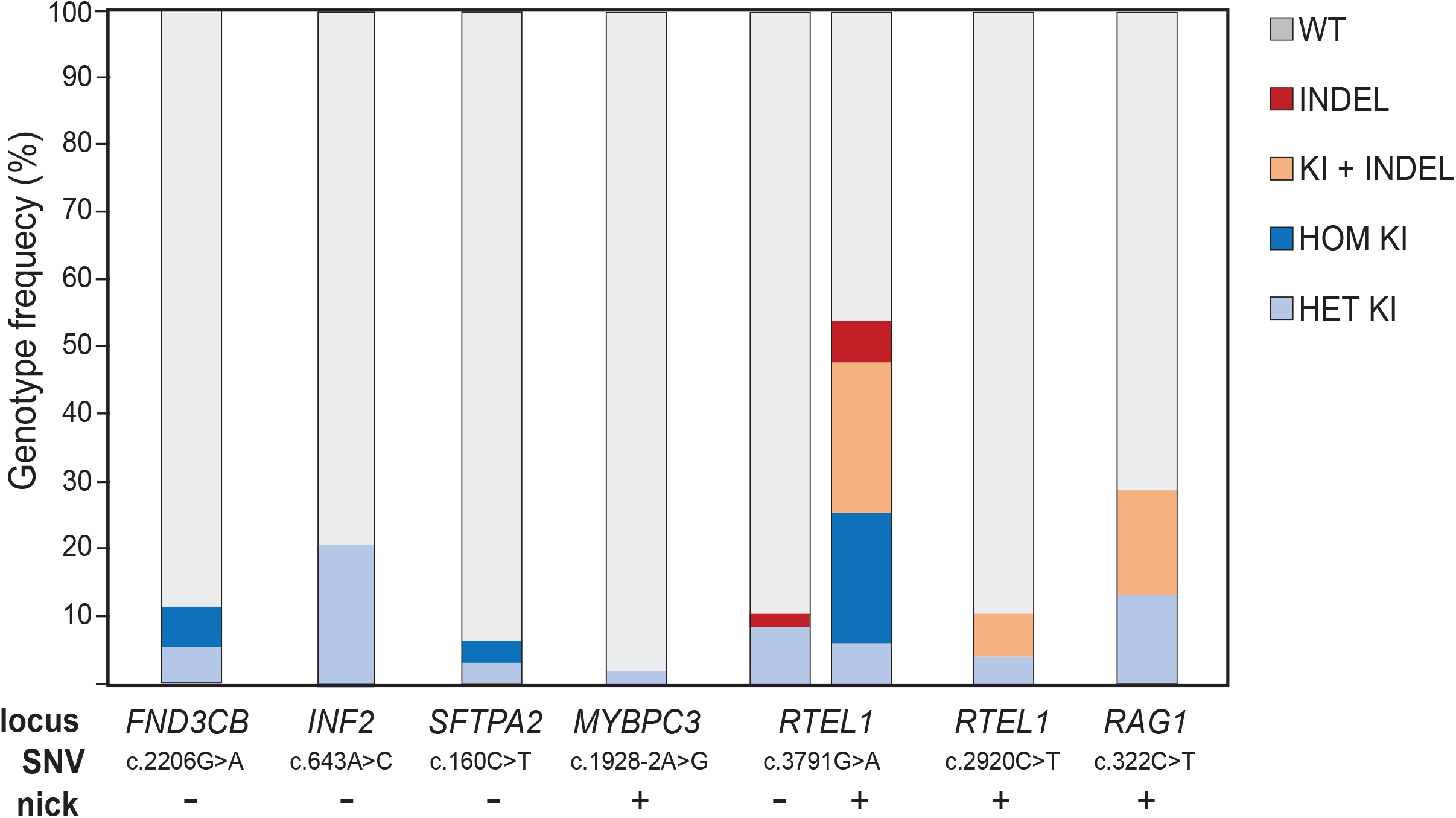
Genotypic analysis of iPSC colonies from further prime-editing experiments. Individual colonies were isolated and analysed by amplicon NGS. Colonies harbouring >50% of the corresponding SNC knock-in allele were classified as homozygous (HOM KI) whereas colonies harbouring 10–50% of the SNV allele were classified as heterozygous (HET KI). Colonies classified as heterozygous knock-in were also confirmed following subclone analysis.

We also sought to evaluate the capacity for PE to generate more complex edits, specifically in loci containing identical sequences within closely related paralogues. The *MYH6* and *MYH7* genes, which encode the alpha and beta isoforms of cardiac myosin heavy chain respectively, share a common evolutionary origin and have both been linked to various human cardiac pathologies.^17^ We attempted to introduce a heterozygous SNV in the *MYH7* locus (c.2155C>T) without affecting corresponding sequences within the *MYH6* locus (c. 2161C>T). Located in tandem, the *MYH6* and *MYH7* genes likely emerged from a previous gene duplication event and share >91% sequence identity^18^, rendering them less amenable to editing strategies that rely on Cas9 endonuclease. Given all potential sgRNAs within an 80 bp window of the *MYH7* c.2155C>T SNV also recognise corresponding sequences in *MYH6*, Cas9 nuclease would likely induce DSBs within both genes, potentially leading to deletion of the intervening (∼30 kb) sequence. We designed two epegRNAs, which encode the C>T SNV as well as spacers that are predicted to bind *MYH7* and *MYH6* with equal efficiency (Figure 3A). To reduce indel formation and minimise multiple editing events we chose to omit a secondary nicking sgRNA. Plasmids encoding either epegRNA-1 or epegRNA-2 along with mRNA encoding PE6b and MLH1dn were introduced into MCRIi010-A iPSCs and editing efficiency was measured in bulk transfected cultures by amplicon NGS, with *MYH6* and *MYH7* alleles distinguishable by two single nucleotide differences located upstream and downstream of the target site (Figure 3A). Despite epegRNA-1 yielding a higher overall editing efficiency, we selected the epegRNA-2 condition for iPSC colony isolation due to a higher ratio of *MYH7* to *MYH6* KI events (Figure 3B). Of the 44 colonies screened, five (11.4%) exhibited SNV knock-in exclusively within *MYH7*, two (4.5%) exclusively within *MYH6*, and six (13.6%) within both *MYH6* and *MYH7* (Figure 3B). All remaining colonies were unedited (≥95% wildtype alleles) and no indel mutations were detected in any of the colonies screened. Next three colonies, harbouring 23–30% *MYH7* c.2155C>T but ≤1% *MYH6* c.2161C>T alleles, were selected for subcloning and subsequent NGS analysis. This enabled successful isolation of multiple iPSC clones from each parent colony, with clean heterozygous knock-in of the *MYH7* c.2155 SNV (as determined by an equal ratio of *MYH7* c.2155C>T to wildtype alleles). Importantly, no subclone showed >50% *MYH7* c.2155C>T alleles indicating homozygous knock-in had not occurred in any of the parent colonies selected for subcloning. SNV knock-in at the *MYH7* locus, but not at *MYH6*, was further confirmed using gene-specific primers to separately amplify *MYH6* and *MYH7*, followed by Sanger sequencing of the resulting amplicons (Figure 3D).

**Figure 3.**
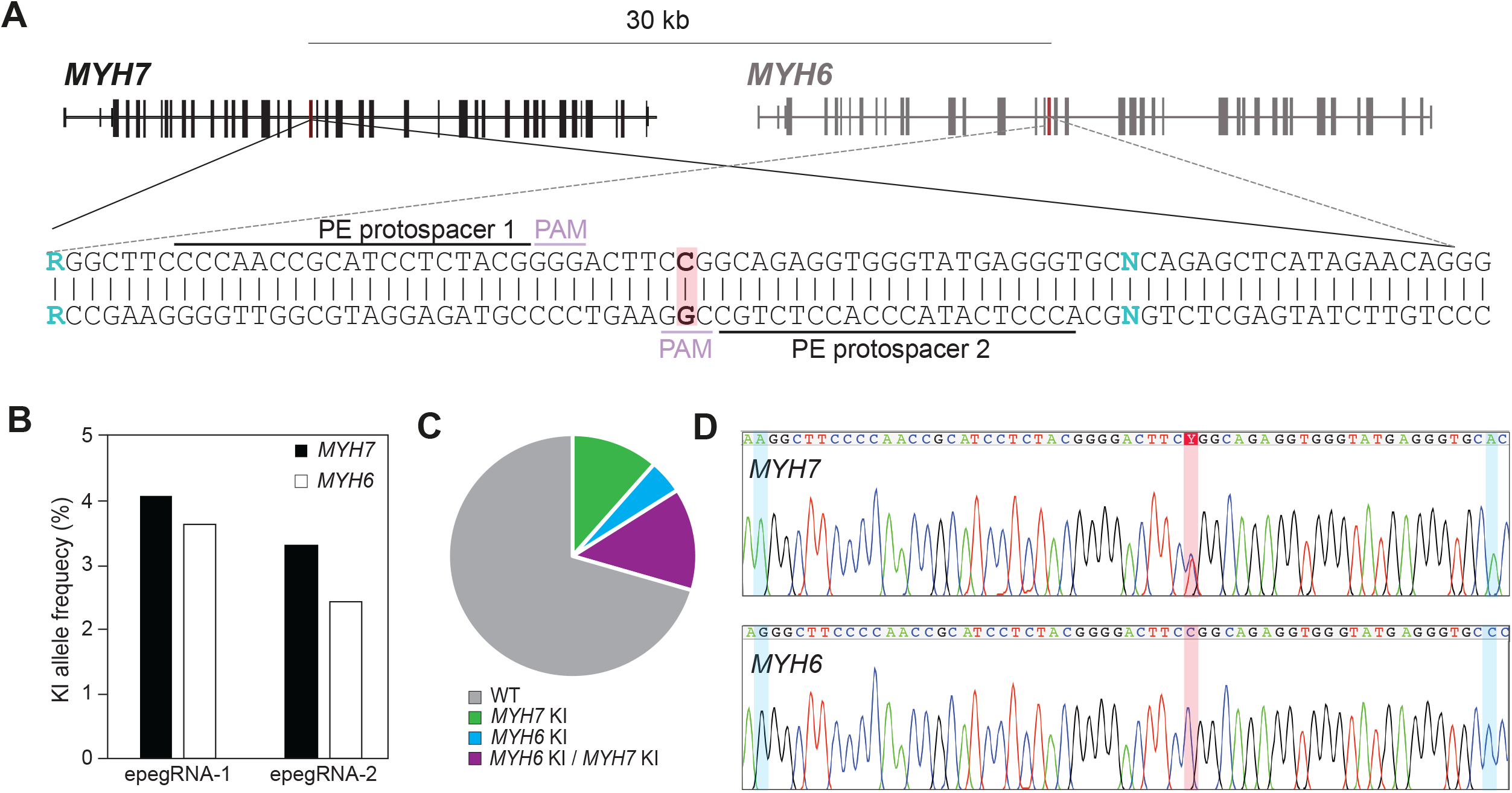
Harnessing PE to install heterozygous edits in paralogous genes. (A) Schematic of the tandemly arranged *MYH7* and *MYH6* genes. The spacers encoded by the two epegRNA constructs, the nucleotide intended for editing (highlighted in red) and the two nucleotides used to distinguish *MYH6* and *MYH7* (blue type) are shown. (B) Percentage of *MYH7* c.2155C>T (black bars) and *MYH6* c.2161C>T alleles (white bars) in bulk transfected iPSC cultures. (C) Proportion of iPSC colonies harbouring knock-in of *MYH7* c.2155C>T, *MYH6* c.2161C>T or both *MYH7* c.2155C>T and *MYH6* c.2161C>T SNVs. (D) Sanger sequencing analysis of a representative iPSC clone harbouring heterozygous knock-in of the *MYH7* c.2155C>T SNV and an unedited *MYH6* locus. The *MYH7* c.2155C>T SNV (highlighted in red) and nucleotides used to distinguish *MYH6* and *MYH7* (highlighted in blue) are shown.

We also utilised PE to introduce a heterozygous SNV (*SFTPA* c.532 G>A) in either the *SFTPA1* or *SFTPA2* genes. *SFTPA1* and *SFTPA2* exhibit >98% sequence homology and have been implicated in adult-onset pulmonary fibrosis and lung cancer.^19,20^ Although in reverse orientation to one another, *SFTPA1* and *SFTPA2* are also arranged in tandem (Figure S4A), making them incompatible with endonuclease-based editing strategies. A nicking sgRNA was included in the PE cocktail along with mRNA encoding PE6b and MLH1dn, since editing efficiency was extremely low (≤0.5%) in its absence. A single nucleotide difference located 74 bp downstream of the *SFTPA* c.532G>A target was used to distinguish *SFTPA1* and *SFTPA2* alleles (Figure S4). Of 72 colonies screened, four (5.6%) exhibited knock-in of the c.532G>A SNV within *SFTPA1* and three (4.2%) within *SFTPA2*. However, two of the *SFTPA2* knock-in clones also harboured a 9 bp deletion – located within the *SFTPA1* gene in one clone and adjacent to the *SFTPA2* c.532G>A SNV in the other - which spanned sequences between the epegRNA and nicking PAM sites. Large deletions exceeding the full length of the NGS amplicon (>200 bp), were also detected in four (4.2%) clones lacking SNV knock-in; two within *SFTPA1* and two in *SFTPA2*. Nonetheless, we subsequently isolated subclones with clean heterozygous incorporation of the c.532G>A mutation exclusively within *SFTPA1* or *SFTPA2*, which was confirmed using gene-specific primers followed by Sanger sequencing of the resulting amplicons (Figure S4C and D).

Lastly, we evaluated the feasibility of utilising PE in our previously described one-step genome editing and reprogramming workflow, which enables the derivation of genetically engineered iPSC clones directly from human fibroblasts.^19,21^ We initially aimed to derive iPSCs from healthy foreskin fibroblasts carrying a heterozygous SNV in *ALPK3* (Figure 4A), a gene that has previously been linked to familial cardiomyopathy.^20^ An epegRNA encoding the *ALPK3* c.1792C>T SNV and an adjacent synonymous change was introduced into primary foreskin fibroblasts (GM21808; Coriell) along with mRNA encoding PEmax in additional to the episomal vectors required for reprogramming. We also performed a direct comparison to our previously established protocol that uses a gene-editing cocktail consisting of a plasmid encoding the gene-specific sgRNA, ssODN repair template and mRNA encoding Cas9-Gem - a Cas9 variant that is associated with lower rates of non-homologous end joining.^13^ In our one-step workflow, iPSC colonies typically emerge from a single fibroblast cell and are therefore clonal in nature, eliminating the need for successive rounds of subcloning. As such, identification of successfully edited iPSC clones can be achieved using basic techniques such as allele-specific PCR and/or Sanger sequencing analysis (Figure 4B and C). In the absence of MLH1dn, we successfully identified one (3.1%) out of 32 iPSC clones harbouring heterozygous incorporation of the *ALPK3* c.1792C>T SNV (Figure 4D). The inclusion of MLH1dn mRNA led to over a fourfold increase in editing efficiency, with six (14%) out of 43 iPSC clones exhibiting heterozygous incorporation of the *ALPK3* c.1792C>T SNV. With respect to the Cas9-Gem mRNA/ssODN condition, we identified two (4.7%) out of 43 iPSC clones harbouring homozygous incorporation of the SNV and two (4.7%) clones with heterozygous SNV knock-in. However, both heterozygous clones also harboured indel mutations (1 bp or 2 bp deletion) in the second *ALPK3* allele. (Figure 4C and D). We also tested whether PE can be used in conjunction with reprogramming to derive gene edited iPSCs directly from patient peripheral blood mononuclear cells (PBMCs) from an infant with a severe form of autosomal recessive polycystic kidney disease (ARPKD) who harbours compound heterozygous mutations in the *PKHD1* gene (Figure S5). We successfully used PE to correct one mutation, *PKHD1* c.107C>T, which is the most common mutation associated with ARPKD.^22^

**Figure 4.**
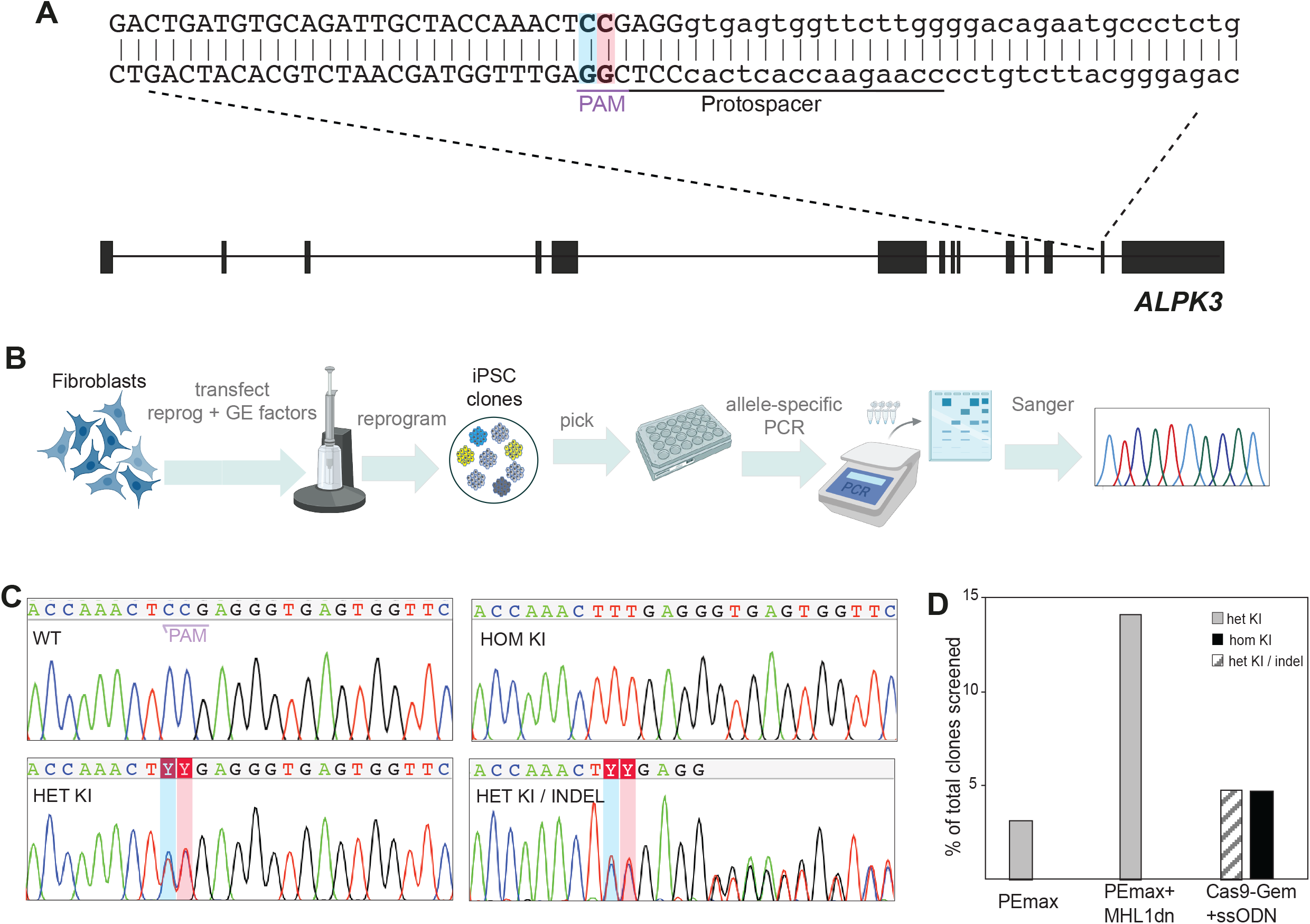
One-step prime-editing and reprogramming of primary fibroblasts. (A) *ALPK3* locus showing location of the *ALPK3* c.1792C>T SNV (highlighted in red) and accompanying synonymous change (highlighted in blue). (B) Overview of the one-step PE/reprogramming workflow used to generate gene-edited iPSCs directly from primary fibroblasts. iPSC clones harbouring knock-in of *ALPK3* c.1792C>T were identified by allele-specific PCR and confirmed by Sanger sequencing (C). (D) Proportion of clones harbouring heterozygous knock-in of the ALPK3 c.1792C>T SNV isolated from PE experiments performed with and without MLH1dn. Only clones with biallelic modifications were identified from the Cas9-Gem mRNA/ssODN condition.

## DISCUSSION

The development of advanced genome engineering systems has profoundly enhanced the potential of hPSCs in developmental biology, disease modelling, and regenerative medicine applications. However, achieving reliable knock-in of SNVs in a heterozygous context can be challenging. Here we comprehensively demonstrate that PE is a robust gene editing tool for installing heterozygous edits in a broad range of genetic loci. We successfully generated iPSC lines carrying 15 unique SNVs across 8 different genetic backgrounds. Moreover, in nearly all instances, we successfully obtained multiple independent clones with heterozygous incorporation of the gene-specific SNV. This was accomplished without the need to screen large numbers of colonies, with no more than 72 individual colonies genotyped per experiment using a workflow that is free of subsequent cell purification or drug selection steps.

We also demonstrate that PE can effectively introduce heterozygous edits in paralogous genes exhibiting very high sequence homology. Paralogues typically arise from duplication events and are generally not suitable for endonuclease-based editing strategies, as dual sgRNA recognition frequently leads to large deletions, inversions, or other chromosomal rearrangements. Indeed, dual sgRNA/Cas9 systems are commonly employed to intentionally induce such outcomes.^23-25^

Consistent with previous reports,^16^ we found the inclusion of a secondary nicking sgRNA resulted in higher editing efficiency, but also led to an increased indel byproducts. Nonetheless, in experiments where we performed direct comparisons with Cas9 endonuclease, PE was associated with a lower overall indel frequency. Going forward, we recommend conducting PE experiments both with and without a nicking sgRNA and evaluating outcomes in bulk transfected cells. If sufficient editing efficiency is achieved without the nicking sgRNA, we suggest proceeding with colony isolation and screening.

While heterozygous knock-in was the main focus of this study, PE also induced homozygous edits, particularly when a secondary nicking sgRNA was included.

The inclusion of mRNA encoding the dominant negative mismatch repair protein, MLH1dn, also consistently improved editing efficiencies. This was particularly evident in the instances where PE was combined with reprogramming to generate gene edited iPSCs directly from primary fibroblasts or PBMCs. Importantly, transient MLH1dn expression was not associated with any genomic aberrations, with no aneuploidies or copy number variations detected in any of the clones generated throughout this study, as determined by SNP array molecular karyotyping analyses.

Taken together, our results demonstrate that PE is a highly effective approach for generating iPSCs with complex genetic modifications, facilitating the study of heterozygous variants within a consistent genetic background. This provides a powerful platform for investigating disease-associated dominant mutations in a fully isogenic context, with the potential to uncover functional outcomes that have yet to be explored.

## METHODS

### Vector construction

PegRNAs-expressing plasmids were generated by ligating two sets of annealed oligo pairs encoding pegRNA scaffold, evopreQ_1_ motif and locus/variant specific the Cas9-target (spacer), PBS and RT sequences was ligated into the BsaI-digested pU6-peg-GG-acceptor plasmid (Addgene #132777; gift from D. Liu). Annealed oligo pairs were treated with T4 DNA kinase, followed by heat inactivation, prior to ligation. Nicking sgRNA plasmids were generated by ligating annealed oligo pairs into BbsI-digested pSMART-sgRNA (Addgene #80427) plasmid. Transfection grade plasmid DNA was prepared using the QIAGEN Plasmid Midi kit. See Table S1 for full list of oligonucleotides used to generate epegRNA and sgRNA constructs.

### In Vitro Transcription

Capped and polyadenylated in vitro transcribed mRNA encoding PEmax and PE6b was generated from PmeI-digested pCMV-PEmax (Addgene #174820; gift from D. Liu) and pCMV-PE6b (Addgene # 207852; gift from D. Liu) plasmid DNA respectively. A gBlock (Integrated DNA Technologies) encoding a T7 promoter and MLH1dn coding sequence was used for in vitro transcription of mRNA encoding MLH1dn. In vitro transcription was performed using the mMESSAGE mMACHINE T7 ULTRA transcription kit (Thermo Fisher) according to the manufacturer’s recommendations. LiCl was used to precipitate mRNA prior to resuspension in nuclease-free H_2_0.

### Amplicon Next Generation Sequencing

Sequencing was performed using Illumina MiSeq technology, following the 16S metagenomic Sequencing Library Preparation protocol for amplicon library preparation and sequencing. Briefly, genomic DNA was extracted from bulk transfected cultures the DNeasy Blood and Tissue Kit (QAGEN) or from isolated iPSC colonies using Quick Extraction Buffer and ∼200 bp amplicons were generated by PCR using adapter primers which flank the target site (see Table S3 for full list of gene-sepcific primers). Exonuclease I (NEB) was then used to catalyse the removal of excess oligonucleotides and amplicons were indexed with dual-barcode primers (Nextera XT DNA Indexes, Illumina). Indexed amplicons underwent fragment analysis and quality control using capillary electrophoresis (QIAxcel connect system, Qiagen) before samples were pooled in equimolar concentrations and cleaned up using Ampure XP magnetic beads (Beckman Coulter) at a 0.8 ratio. The pooled library was denatured and loaded onto the MiSeq sequencing platform with a final concentration of 6 pM + 15% PhiX spike-in and 300 bp paired-end sequencing was performed (MiSeq Reagent Nano Kit v2 300 cycles, Illumina). All PCR amplifications were carried out using Q5 High-Fidelity DNA polymerase (NEB).

Sequencing reads were aligned to the GRCh38 reference genome using “bwa mem” tool. Reads were reformatted with “samtools” to BAM files. A custom R script using the “CrispRVariants” package^26^ was used to evaluate and plot gene editing outcomes.Briefly, using the “readsToTarget ()” function, reads aligned to the target site were extracted from the BAM files, producing a “CrisprSet” object. Sequencing variants near the target location were quantified and then visualised using the “plotVariants()” function, generating heat maps of variant frequencies across samples. Coverage at target SNV sites averaged at ∼8,000 reads and ranged from 5,000 to 11,000 reads. All NGS data is available through Mendeley Data (DOI: 10.17632/yn7kpg48ym.1) in agreement with FAIR principles. https://data.mendeley.com/datasets/yn7kpg48ym/1

### PCR and Sanger Sequencing

Allele-specific PCR was performed using GoTaq Green PCR Mastermix (Promega) using primers depicted in Table S4. Prior to sequencing, PCR amplicons were treated with rAPid Alkaline Phosphatase (Roche Life Science) and Endonuclease I (NEB) for 30 min at 37°C followed by heat inactivation at 80°C for 15 min. Sequencing reactions were performed using BigDye Terminator v3.1 (Thermo Fisher Scientific). Reactions were purified and sequenced by the Australian Genome Research Facility.

### Cell culture, cryopreservation and quality assurance

The iPSC lines used in this study are listed in Table S4. Generation and characterisation of PB010.5, PB005.1 and PB001.1 iPSC lines have been described previously.^27^ iPSCs were maintained and expanded at 37°C, 5% CO_2_ and 20% O_2_ in Essential 8 (Thermo Fisher Scientific) or mTesR+ media (Stem Cell Technologies) on hESC-qualified Matrigel (Corning) coated plates with daily media changes. No antibiotics were used in the culture medium. iPSCs were passaged using 0.5 mM EDTA in 1X PBS every 3–4 days when cultures reached approximately 80% confluency and/or when individual colonies 1–2 mm in size as previously described.^28^ For colony picking, single iPSC colonies (approximately 1–2 mm in size) were detached with a sterile 200 μl filter tip and transferred to one well of 48-well Matrigel-coated plate. All SNV knock-in experiments described in this study were requested by users of the iPSC Gene Editing Core Facility (Murdoch Children’s Research Institute, Australia) and iPSC clones confirmed to harbour heterozygous knock-in of the requested variant were subjected to standard quality assurance procedures (molecular karyotyping, assessment of pluripotency markers, mycoplasma testing) after expansion and cryopreservation but prior to handover. Molecular karyotyping was performed by Victorian Clinical Genetics Services (Parkville, Australia), using Illumina Infinium CoreExome-24 v3.0 SNP arrays. SNPduo analyses were performed to confirm gene-edited clones were derived from the corresponding parental iPSC line. For assessment of pluripotency markers, iPSCs were harvested with TrypLE (Thermo Fisher Scientific) and incubated with antibodies to cell surface markers TRA-1-60 BV421 (BD Horizon), TRA-1-81 AF647 (Biolegend) and SSEA4 PeCy7 (Biolegend) as per manufacturer’s specifications. For intracellular OCT4 staining, cells were treated with Foxp3 fixation/permeabilisation buffer (Thermo Fisher Scientific) and stained with OCT3/4 PE antibody (Thermo Fisher Scientific). Cells were analysed by flow cytometry using a LSRFortessa x-20 flow cytometer (BD Biosciences). Unstained iPSCs were used as a gating control. Mycoplasma testing was performed by Cerberus Sciences (Scoresby, Australia). For cryopreservation, iPSCs were dissociated 2 days post passaging (at approximately 50% confluency) using 0.5 mM EDTA in 1x PBS and resuspended in Essential 8 medium. An equal volume of freezing medium (20% DSMO in Essential 8 medium) was added dropwise and the cell suspension mixed by gentle agitation prior to aliquoting (1–1.5 ml) into cyrotubes for controlled rate freezing. Thawed cells were transferred to 15 ml Falcon tubes and 5 ml Essential medium added dropwise to the cell suspension prior to centrifugation (300g for 3 min). The cell pellet was resuspended in 2 ml Essential 8 medium and plated at varying densities across two wells of a Matrigel-coated 24-well plate.

### Subcloning

Subcloning was performed by dissociating iPSC cultures 1–2 days after EDTA passaging with TrypLE. Cells were plated at low density (100–1000 cells per well) in 6-well Matrigel-coated plates in Essential 8 medium supplemented with CloneR2 (Stem Cell Technologies). The next day, colonies consisting of 2–3 cells were marked and allowed to expand for ∼ 1 week prior to picking, expansion and subsequent NGS analysis.

### Transfection

iPSC transfections were performed using the Neon Transfection System (Thermo Fisher Scientific) as previously described.^13^ Briefly, cells were dissociated with TrypLE and resuspended in Buffer R at a final concentration of 1 × 10^7^ cells/ml. Transfections were performed in 100 μl tips using the following conditions: 1100 V, 30 ms, one pulse. Following transfection, cells were transferred to Matrigel-coated plates containing mTesR+ medium supplemented with CloneR2, which was omitted in subsequent media changes. Simultaneous reprogramming and gene editing of human fibroblasts (GM21808, Coriell) was performed as previously described.^19^ Briefly, fibroblasts were dissociated with TrypLE two days after passaging and resuspended in Buffer R at a final concentration of 1 × 10^7^ cells/ml. Electroporation was performed in 100 μl tips using the Neon Transfection device using the following conditions: 1,400 V, 20 ms, two pulses. Cells were subsequently seeded on Matrigel-coated plates and maintained in fibroblast medium (DMEM supplemented with 10% fetal bovine serum) until three days post transfection, then switched to E7 medium (E8 medium without transforming growth factor β) supplemented with 100 μM sodium butyrate and changed every other day. Sodium butyrate was removed from the medium after the appearance of the first iPSC colonies at around day 10.

## AUTHOR CONTRIBUTIONS

S.E.H. designed experiments and wrote the manuscript. A.S performed NGS experiments with assistance from J.K. and J.C. J.C and A.S constructed, prepped and sequence verified all epegRNA and sgRNA constructs. A.S performed NGS analysis. A.S, J.K. K.V, A.G, K.S, and C.G all provided assistance with cell maintenance, transfection, cryopreservation, colony picking and quality assurance. F.K.R and M.S. and M.R provided bioinformatics support including set up of the custom CrispRVariants pipeline.

## ACKNOWLEDGEMENTS

We thank users of the MCRI iPSC Gene Editing Facility for allowing us to share information around the generation of their requested iPSC lines. The iPSC Gene Editing Facility was established using a generous donation from the Stafford Fox Medical Research Foundation and is currently supported by Phenomics Australia (PA), and the Novo Nordisk Foundation reNEW Center for Stem Cell Medicine (NNF21CC0073729). PA is supported by the Australian Government through the National Collaborative Research Infrastructure Strategy program.

## REFERENCES

1. Dorison, A., Ghobrial, I., Graham, A., Peiris, T., Forbes, T.A., See, M., Das, M., Saleem, M.A., Quinlan, C., Lawlor, K.T., et al. (2023). Kidney Organoids Generated Using an Allelic Series of NPHS2 Point Variants Reveal Distinct Intracellular Podocin Mistrafficking. J Am Soc Nephrol 34, 88–109. 10.1681/asn.2022060707.

2. Cong, L., Ran, F.A., Cox, D., Lin, S., Barretto, R., Habib, N., Hsu, P.D., Wu, X., Jiang, W., Marraffini, L.A., and Zhang, F. (2013). Multiplex genome engineering using CRISPR/Cas systems. Science 339, 819–823. 10.1126/science.1231143.

3. Mali, P., Yang, L., Esvelt, K.M., Aach, J., Guell, M., DiCarlo, J.E., Norville, J.E., and Church, G.M. (2013). RNA-guided human genome engineering via Cas9. Science 339, 823–826. 10.1126/science.1232033.

4. Jinek, M., East, A., Cheng, A., Lin, S., Ma, E., and Doudna, J. (2013). RNA-programmed genome editing in human cells. Elife 2, e00471. 10.7554/eLife.00471.

5. Anzalone, A.V., Randolph, P.B., Davis, J.R., Sousa, A.A., Koblan, L.W., Levy, J.M., Chen, P.J., Wilson, C., Newby, G.A., Raguram, A., and Liu, D.R. (2019). Search-and-replace genome editing without double-strand breaks or donor DNA. Nature 576, 149–157. 10.1038/s41586-019-1711-4.

6. Kim, D.Y., Moon, S.B., Ko, J.H., Kim, Y.S., and Kim, D. (2020). Unbiased investigation of specificities of prime editing systems in human cells. Nucleic Acids Res 48, 10576–10589. 10.1093/nar/gkaa764.

7. Li, H., Busquets, O., Verma, Y., Syed, K.M., Kutnowski, N., Pangilinan, G.R., Gilbert, L.A., Bateup, H.S., Rio, D.C., Hockemeyer, D., and Soldner, F. (2022). Highly efficient generation of isogenic pluripotent stem cell models using prime editing. Elife 11. 10.7554/eLife.79208.

8. Li, M., Zhong, A., Wu, Y., Sidharta, M., Beaury, M., Zhao, X., Studer, L., and Zhou, T. (2022). Transient inhibition of p53 enhances prime editing and cytosine base-editing efficiencies in human pluripotent stem cells. Nat Commun 13, 6354. 10.1038/s41467-022-34045-7.

9. Chemello, F., Chai, A.C., Li, H., Rodriguez-Caycedo, C., Sanchez-Ortiz, E., Atmanli, A., Mireault, A.A., Liu, N., Bassel-Duby, R., and Olson, E.N. (2021). Precise correction of Duchenne muscular dystrophy exon deletion mutations by base and prime editing. Sci Adv 7. 10.1126/sciadv.abg4910.

10. Sürün, D., Schneider, A., Mircetic, J., Neumann, K., Lansing, F., Paszkowski-Rogacz, M., Hänchen, V., Lee-Kirsch, M.A., and Buchholz, F. (2020). Efficient Generation and Correction of Mutations in Human iPS Cells Utilizing mRNAs of CRISPR Base Editors and Prime Editors. Genes (Basel) 11. 10.3390/genes11050511.

11. Herman, D.S., Lam, L., Taylor, M.R., Wang, L., Teekakirikul, P., Christodoulou, D., Conner, L., DePalma, S.R., McDonough, B., Sparks, E., et al. (2012). Truncations of titin causing dilated cardiomyopathy. N Engl J Med 366, 619–628. 10.1056/NEJMoa1110186.

12. Nelson, J.W., Randolph, P.B., Shen, S.P., Everette, K.A., Chen, P.J., Anzalone, A.V., An, M., Newby, G.A., Chen, J.C., Hsu, A., and Liu, D.R. (2022). Engineered pegRNAs improve prime editing efficiency. Nat Biotechnol 40, 402–410. 10.1038/s41587-021-01039-7.

13. Howden, S.E., McColl, B., Glaser, A., Vadolas, J., Petrou, S., Little, M.H., Elefanty, A.G., and Stanley, E.G. (2016). A Cas9 Variant for Efficient Generation of Indel-Free Knockin or Gene-Corrected Human Pluripotent Stem Cells. Stem Cell Reports 7, 508–517. 10.1016/j.stemcr.2016.07.001.

14. Bonnycastle, L.L., Swift, A.J., Mansell, E.C., Lee, A., Winnicki, E., Li, E.S., Robertson, C.C., Parsons, V.A., Huynh, T., Krilow, C., et al. (2024). Generation of Human Isogenic Induced Pluripotent Stem Cell Lines with CRISPR Prime Editing. Crispr j 7, 53–67. 10.1089/crispr.2023.0066.

15. Doman, J.L., Pandey, S., Neugebauer, M.E., An, M., Davis, J.R., Randolph, P.B., McElroy, A., Gao, X.D., Raguram, A., Richter, M.F., et al. (2023). Phage-assisted evolution and protein engineering yield compact, efficient prime editors. Cell 186, 3983–4002.e3926. 10.1016/j.cell.2023.07.039.

16. Doman, J.L., Sousa, A.A., Randolph, P.B., Chen, P.J., and Liu, D.R. (2022). Designing and executing prime editing experiments in mammalian cells. Nat Protoc 17, 2431–2468. 10.1038/s41596-022-00724-4.

17. Marian, A.J., and Braunwald, E. (2017). Hypertrophic Cardiomyopathy: Genetics, Pathogenesis, Clinical Manifestations, Diagnosis, and Therapy. Circ Res 121, 749–770. 10.1161/circresaha.117.311059.

18. McGuigan, K., Phillips, P.C., and Postlethwait, J.H. (2004). Evolution of sarcomeric myosin heavy chain genes: evidence from fish. Mol Biol Evol 21, 1042–1056. 10.1093/molbev/msh103.

19. Howden, S.E., Thomson, J.A., and Little, M.H. (2018). Simultaneous reprogramming and gene editing of human fibroblasts. Nat Protoc 13, 875–898. 10.1038/nprot.2018.007.

20. Legendre, M., Butt, A., Borie, R., Debray, M.P., Bouvry, D., Filhol-Blin, E., Desroziers, T., Nau, V., Copin, B., Dastot-Le Moal, F., et al. (2020). Functional assessment and phenotypic heterogeneity of SFTPA1 and SFTPA2 mutations in interstitial lung diseases and lung cancer. Eur Respir J 56. 10.1183/13993003.02806-2020.

21. Howden, S.E., Maufort, J.P., Duffin, B.M., Elefanty, A.G., Stanley, E.G., and Thomson, J.A. (2015). Simultaneous Reprogramming and Gene Correction of Patient Fibroblasts. Stem Cell Reports 5, 1109–1118. 10.1016/j.stemcr.2015.10.009.

22. Goggolidou, P., and Richards, T. (2022). The genetics of Autosomal Recessive Polycystic Kidney Disease (ARPKD). Biochim Biophys Acta Mol Basis Dis 1868, 166348. 10.1016/j.bbadis.2022.166348.

23. Blasco, R.B., Karaca, E., Ambrogio, C., Cheong, T.C., Karayol, E., Minero, V.G., Voena, C., and Chiarle, R. (2014). Simple and rapid in vivo generation of chromosomal rearrangements using CRISPR/Cas9 technology. Cell Rep 9, 1219–1227. 10.1016/j.celrep.2014.10.051.

24. Chen, X., Xu, F., Zhu, C., Ji, J., Zhou, X., Feng, X., and Guang, S. (2014). Dual sgRNA-directed gene knockout using CRISPR/Cas9 technology in Caenorhabditis elegans. Sci Rep 4, 7581. 10.1038/srep07581.

25. Choi, P.S., and Meyerson, M. (2014). Targeted genomic rearrangements using CRISPR/Cas technology. Nat Commun 5, 3728. 10.1038/ncomms4728.

26. Lindsay, H., Burger, A., Biyong, B., Felker, A., Hess, C., Zaugg, J., Chiavacci, E., Anders, C., Jinek, M., Mosimann, C., and Robinson, M.D. (2016). CrispRVariants charts the mutation spectrum of genome engineering experiments. Nat Biotechnol 34, 701–702. 10.1038/nbt.3628.

27. Vlahos, K., Sourris, K., Mayberry, R., McDonald, P., Bruveris, F.F., Schiesser, J.V., Bozaoglu, K., Lockhart, P.J., Stanley, E.G., and Elefanty, A.G. (2019). Generation of iPSC lines from peripheral blood mononuclear cells from 5 healthy adults. Stem Cell Res 34, 101380. 10.1016/j.scr.2018.101380.

28. Chen, G., Gulbranson, D.R., Hou, Z., Bolin, J.M., Ruotti, V., Probasco, M.D., Smuga-Otto, K., Howden, S.E., Diol, N.R., Propson, N.E., et al. (2011). Chemically defined conditions for human iPSC derivation and culture. Nat Methods 8, 424–429. 10.1038/nmeth.1593.

