## Supplemental Figures for "Efficient installation of heterozygous mutations in human pluripotent stem cells using prime editing"

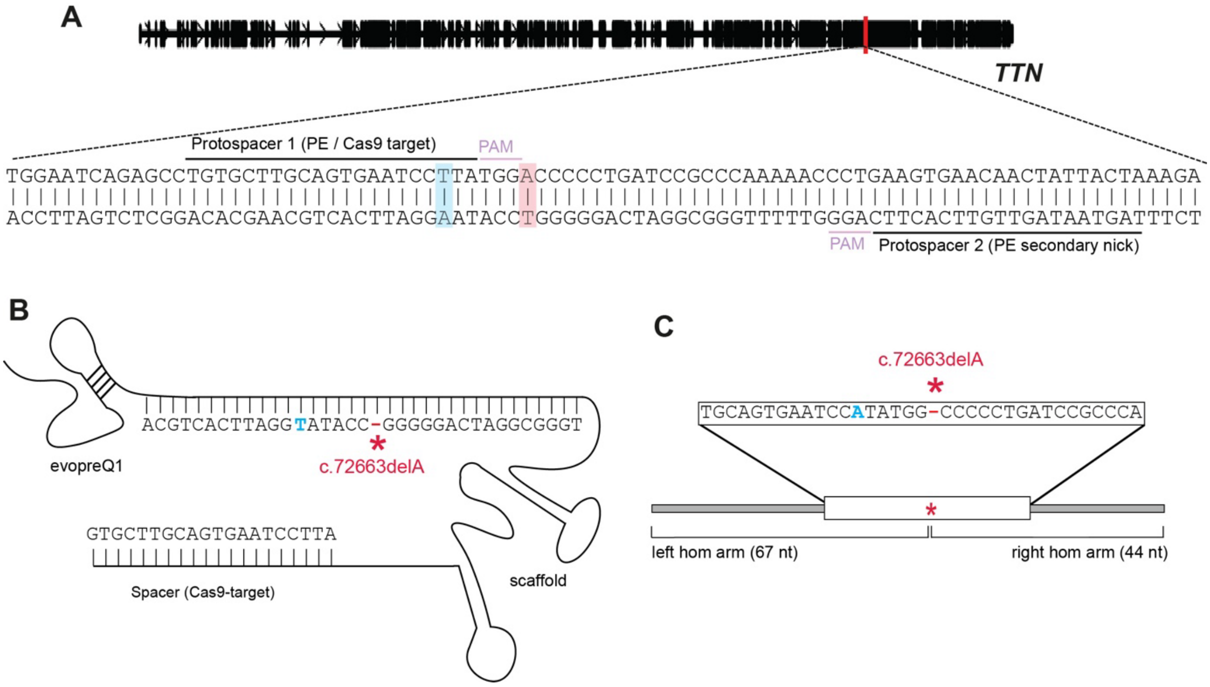


**Supplementary Figure 1. Strategy used to introduce a *TTN* c.72663delA truncating mutation in wildtype iPSCs.** (A) *TTN* locus showing location of the target site. Protospacer 1 is encoded by the epegRNA and sgRNA used in RNP experiments. Protospacer 2 is encoded by the nicking sgRNA, used in a subset of PE experiments. Schematic diagram of the epegRNA (A) and ssODN repair template (B) used in conjunction with RNP. Both encode the *TTN* c.72663delA mutation and a nearby synonymous change (*TTN* c.72657T>A).

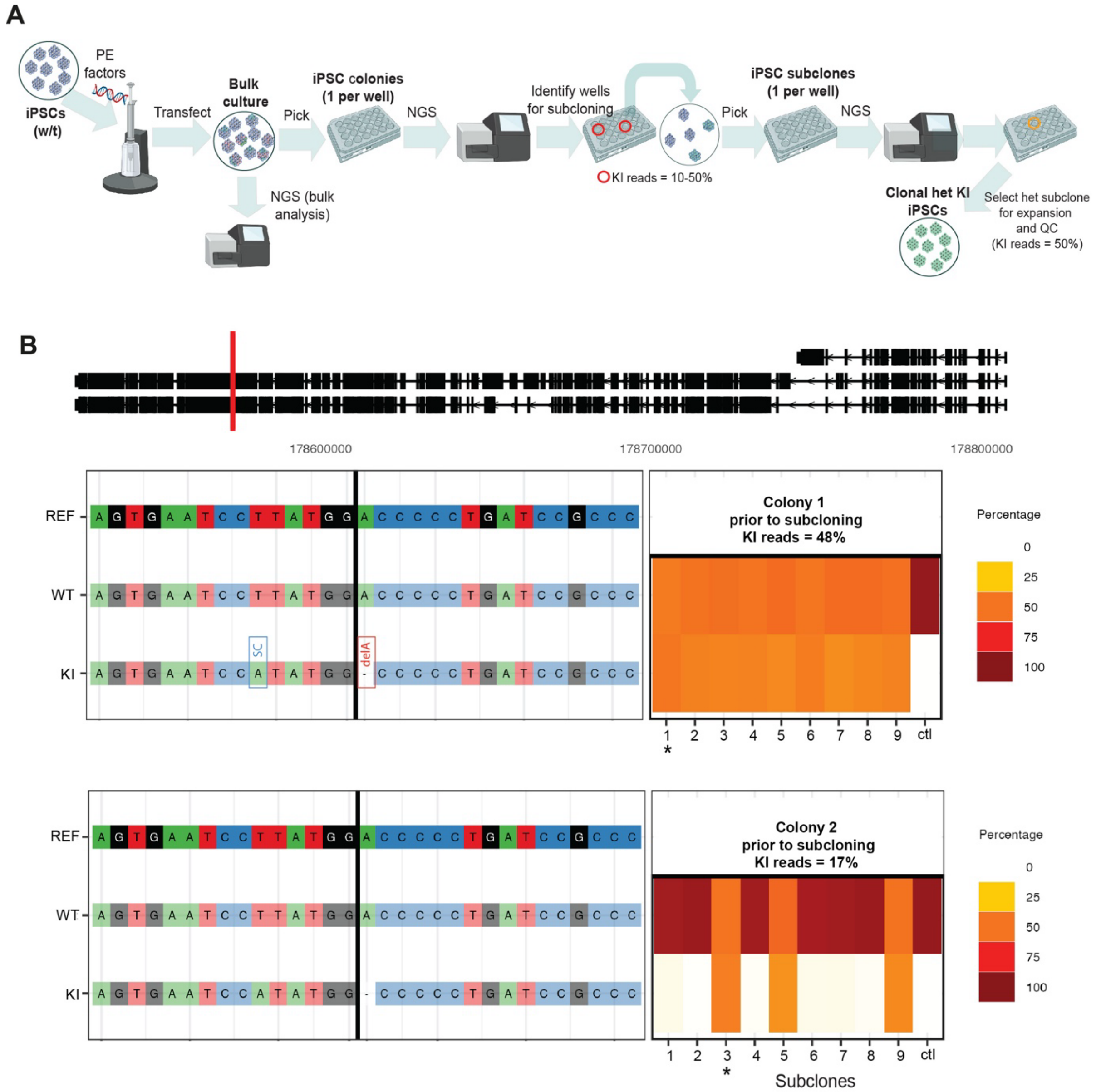


**Supplementary Figure 2. Overview of workflow used to install heterozygous edits and isolate clonal heterozygous iPSC populations.** (A) Gene editing factors are introduced into iPSCs via electroporation. Individual colonies are isolated, expanded and screened by amplicon NGS. Colonies that contain 10-50% knock-in (KI) alleles/reads and therefore likely to contain iPSCs harbouring heterozygous edits are selected for subcloning to attain clonal iPSC cultures. At least 10 subclones are analysed by NGS amplicon sequencing and subclones with a 1:1 ratio of KI:wildtype alleles are selected for expansion and QC. (B) Representative NGS analysis of subclones derived from two parent colonies initially harbouring 48% and 17% *TTN* c.72663delA ; c.72657T>A KI alleles. Subclones (marked with *) confirmed to harbour heterozygous KI (1:1 ratio of KI:wildtype alleles) were selected for expansion and QC.

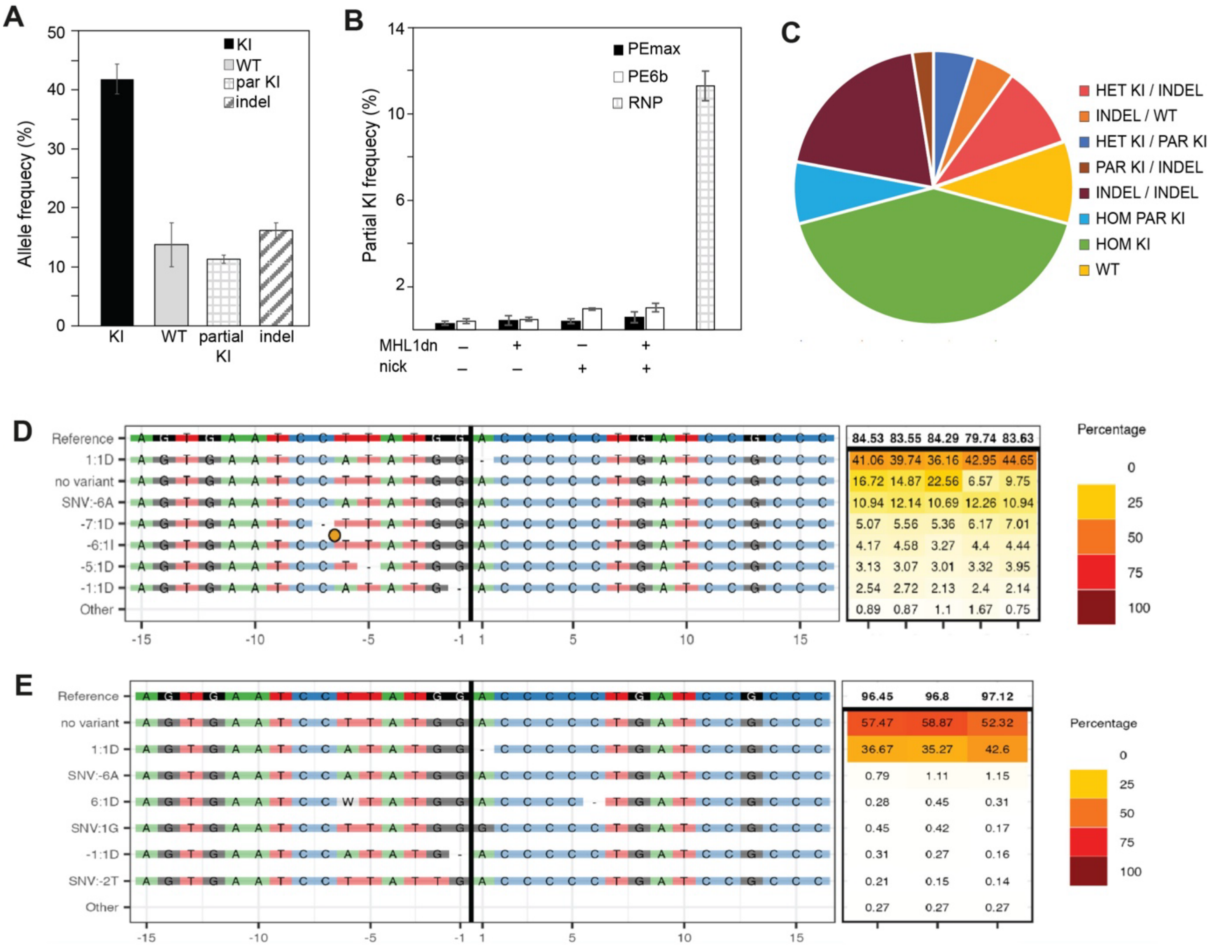


**Supplementary Figure 3. RNP-mediated knock-in of TTN c.72663delA mutation in iPSCs.** (A) Evaluation of editing outcomes, measured as total allele frequency, in bulk iPSC cultures following introduction of *TTN*-specific RNP complex and repair template (a single-stranded oligonucleotide encoding TTN c.72663delA mutation and TTN c.72657T>A synonymous change). N=5. Error bars indicate SD, N=number of biological replicates). (B) Direct comparison of partial KI frequency (incorporation of the *TTN* c.72657T>A synonymous change but not the *TTN* c.72663delA mutation) following introduction of RNP or PE factors in iPSCs. (C) Genotypic analysis of individual colonies isolated from RNP transfection experiments. (D) CrispRVariants plot showing frequency of KI and indel alleles in bulk transfected iPSC cultures following introduction of *TTN*-specific RNP and repair template. N=5. (E) CrispRVariants plot showing frequency of KI and indel alleles in bulk transfected iPSC cultures following introduction of PE factors ("PE6b + MHL1dn + nick" condition). N=3.

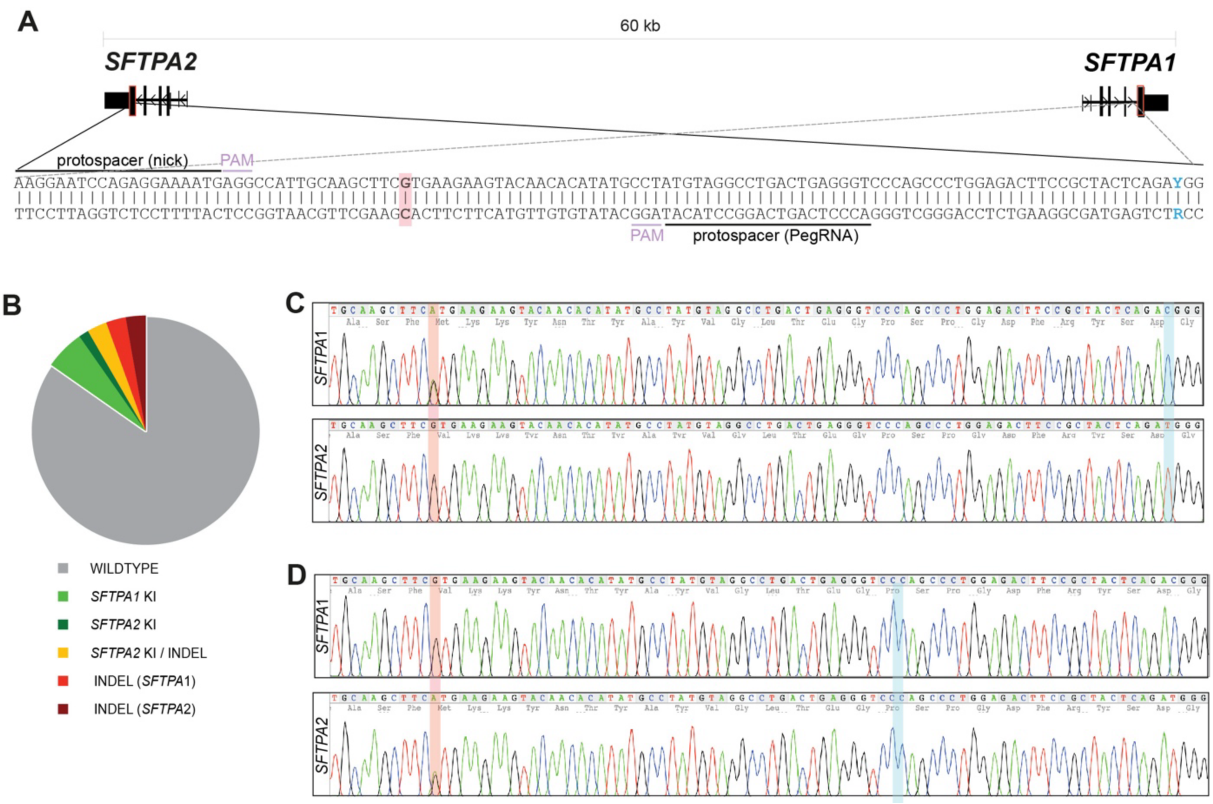


**Supplementary Figure 4. Harnessing prime editing to install heterozygous edits in the *SFTPA* locus**. (A) Schematic of the *SFTPA* locus, consisting of the *SFTPA1* and *SFTPA2* genes which are in reverse transcriptional orientation to one another. Location of the PE and nicking protospacers, the nucleotide intended for editing (highlighted in red) and the single nucleotide used to distinguish *SFTPA1* and *SFTPA2* (highlighted in blue) are shown. (B) Proportion of iPSC colonies harbouring knock-in of *SFTPA1* c.532G>A, *SFTPA1* c.532G>A and/or indel mutations. Sanger sequencing analysis of a representative iPSC clone harbouring clean heterozygous knock-in of *SFTPA1* c.532G>A (C) or *SFTPA2* c.532G>A (D).

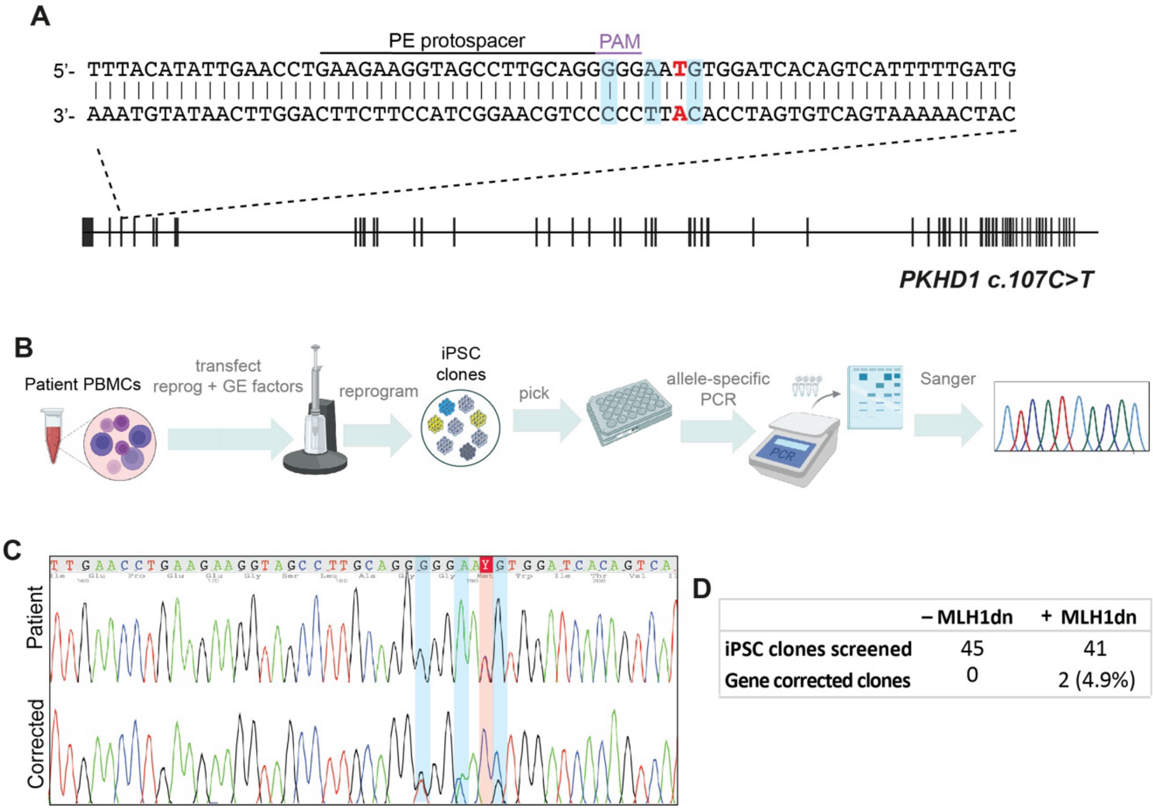


**Supplementary Figure 5. One-step prime-editing and reprogramming of patient PBMCs.** (A) *PKHD1* locus showing location of the *PKHD1* c.107C>T mutation (highlighted in red) and synonymous changes (highlighted in blue) used to identify successfully edited clones. (B) Overview of the one-step PE/reprogramming workflow used to generate gene-edited iPSCs directly from PBMCs. Gene-corrected iPSC clones were identified by allele-specific PCR and confirmed by Sanger sequencing analysis (C). (D) Number of gene-corrected iPSC clones isolated from PE experiments performed with and without MLH1dn.

**Table S4**. iPSC lines used in this study

| Cell line (hPSCreg ID / vendor) | Source | Sex | Culture medium | Passage no. (prior to transfection) | Passage no. (post modification and QC) |
|---|---|---|---|---|---|
| PB010.5 (MCRIi005-A) | PBMCs | Male | Essential 8 | 17-20 | 25-30 |
| PB005.1 (MCRIi005-A) | PBMCs | Female | Essential 8 | 18-20 | 25-27 |
| PB001.1 (MCRIi001-A) | PBMCs | Male | Essential 8 | 17-20 | 25-30 |
| ChiPSC18 (Cellartis, Cat#Y00300) | Fibroblasts | Male | mTesR+ | 18-21 | 27-32 |
